# Treated wastewater irrigation promotes the spread of antibiotic resistance into subsoil pore-water

**DOI:** 10.1101/2020.07.27.222497

**Authors:** Ioannis D. Kampouris, Uli Klümper, Damiano Cacace, Steffen Kunze, Thomas U. Berendonk

## Abstract

In the present study, we investigated the impact of treated wastewater (TWW) irrigation on the prevalence of antibiotic resistance genes (ARGs) in subsoil pore-water, a so-far under-appreciated matrix. We hypothesized that TWW irrigation increases ARG prevalence in subsoil pore-water. This hypothesis was tested using a multiphase approach, which consisted of sampling percolated subsoil pore-water from lysimeter-wells of a real-scale TWW-irrigated field, operated for commercial farming practices, and controlled, laboratory mesocosms irrigated with freshwater or TWW. We monitored the abundance of six selected ARGs (*sul1, bla*_OXA-58_, *tetM*, *qnrS*, *bla*_CTX-M-32_ and *bla*_TEM_), the *intI1* gene associated with mobile genetic elements and an indicator for anthropogenic pollution and bacterial abundance (16S rRNA gene) by qPCR. The bacterial load of subsoil pore water was independent of both, irrigation intensity in the field study and irrigation water type in the mesocosms. Among the tested genes in the field study, *sul1* and *intI1* exhibited constantly higher relative abundances. Their abundance was further positively correlated with increasing irrigation intensity. Controlled mesocosm experiments verified the observed field study results: the relative abundance of several genes, including *sul1* and *intI1,* increased significantly when irrigating with TWW compared to freshwater irrigation. Overall, TWW irrigation promoted the spread of ARGs and *intI1* in the subsoil pore-water, while the bacterial load was maintained. The combined results from the real-scale agricultural field and the controlled lab mesocosms indicate that the dissemination of ARGs in various subsurface environments needs to be taken into account during TWW irrigation scenarios.

**Graphical abstract:** 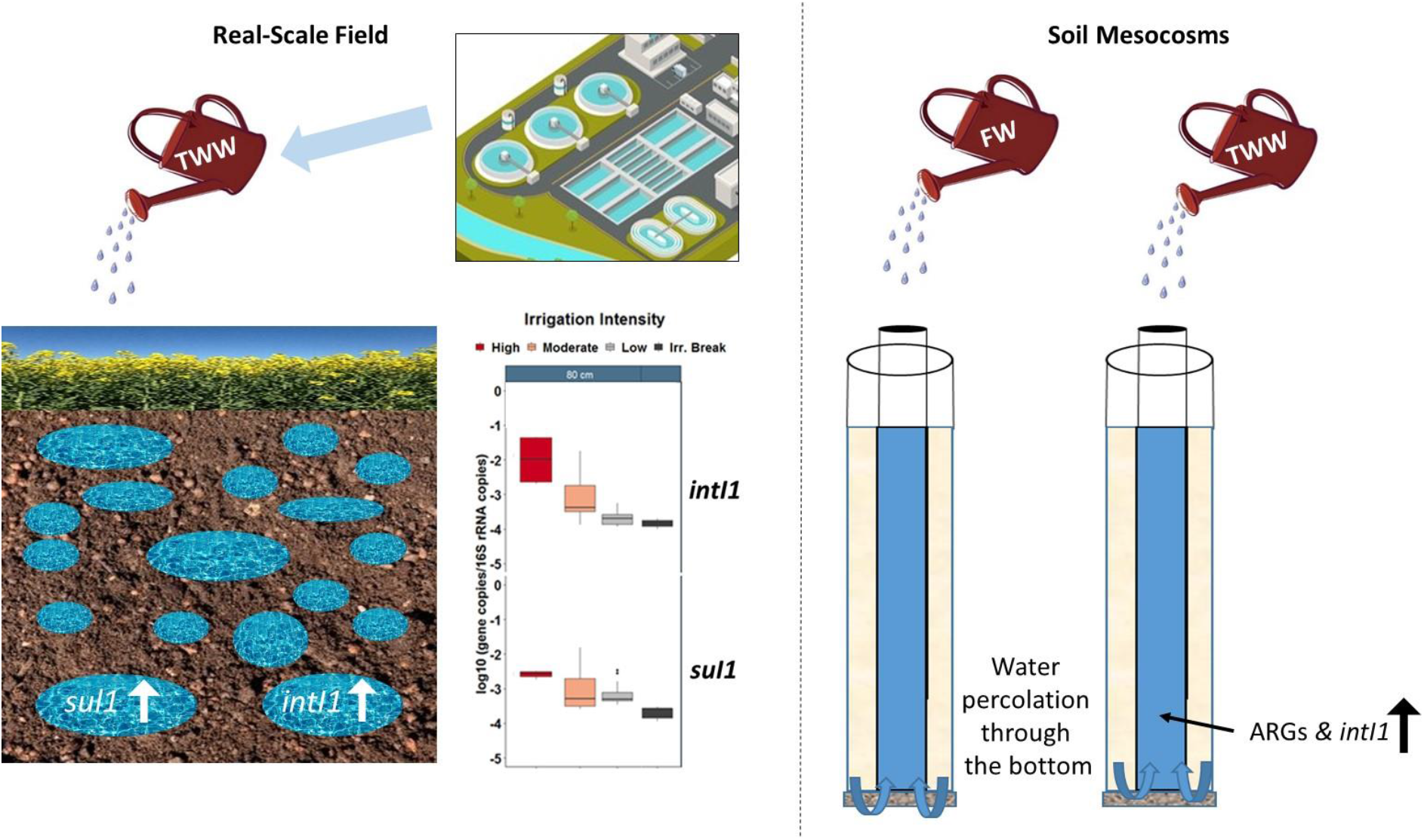

**Highlights:**

- TWW irrigation intensity and *sul1* & *intI1* abundance correlate in a real-scale field
- ARGs & *intI1* increase in subsoil pore-water during TWW irrigation in mesocosms
- No increase of ARGs & *intI1* in freshwater irrigated mesocosms
- TWW irrigation does not affect the bacterial load of subsoil pore-water

## 1. Introduction

Treated wastewater (TWW) irrigation and managed aquifer recharge (MAR) are cost-efficient counter-measures against freshwater resource depletion in arid and semi-arid areas (Paranychianakis et al., 2015; Ternes et al., 2007; Maaß & Grundmann, 2016). However, TWW might contain diverse antibiotic residues (As), antibiotic resistant bacteria (ARB) and antibiotic resistance genes (ARGs) (Berendonk et al., 2015; Cacace et al., 2019; Manaia et al., 2018a; Michael et al., 2013; Pärnänen et al., 2019; Smalla et al., 2018). The abundance of As, ARB and ARGs in TWW has raised questions regarding its influence on the dissemination of antibiotic resistance through irrigation (Michael et al., 2013; Berendonk et al., 2015; Manaia et al., 2018a; Smalla et al., 2018). Previous studies from various locations (China, Australia, Israel, Spain) reported contradictive observations regarding the impact of TWW irrigation on the prevalence of ARGs in soil (Negreanu et al., 2012; Cerquiera et al., 2019a; Cerquiera et al., 2019b; Cerquiera et al., 2019c; Marano et al., 2019; Wang et al., 2014; Han et al., 2016; Dalkmann et al., 2012; Jechalke et al., 2015). Specifically, a few studies have claimed that TWW irrigation increases ARG prevalence in soil (Wang et al., 2014; Han et al., 2016; Dalkmann et al., 2012; Jechalke et al., 2015), while others from the Mediterranean Basin reported a minimal and negligible impact (Negreanu et al., 2012; Cerquiera et al., 2019b; Cerquiera et al., 2019c; Marano et al., 2019). These contradictions could be caused by several factors, such as selection of investigated ARGs, masking from soil-amendment practices, variability of soil microbiota or different A/ARB/ARG-loads present in TWW (Cacace et al., 2019; Pärnänen et al., 2019; Cerquiera et al., 2019c; Marano et al., 2019; Wang et al., 2014; Han et al., 2016; Dalkmann et al., 2012; Jechalke et al., 2015). In addition, effects of irrigation could be influenced by the initial ARG prevalence of the soil microbiome caused by either soil being a natural reservoir of ARGs (native soil resistome), due to the presence of antibiotic producing soil fungi or bacteria (Nesme & Simonet, 2015) or previous exposure to manure (Luby et al., 2016; Yang et al., 2016; Barrios et al., 2020) or biosolids (Mathews & Reinhold, 2013).

Apart from immediate effects on soil, anthropogenic processes can potentially affect the subsoil and groundwater microbiota as well (Szekeres et al., 2018; Rossi et al., 2019). Specifically, Szekeres et al. (2018) reported higher relative abundance of ARGs in groundwater wells with higher proximity to urban settings, due to general anthropogenic activity. However, the majority of studies so far have focused on either crops or topsoil, hence neglecting deeper lying matrices like subsoil, subsoil pore-water and groundwater (Negreanu et al., 2012; Cerquiera et al., 2019a; Cerquiera et al., 2019b; Cerquiera et al., 2019c; Marano et al., 2019; Wang et al., 2014; Han et al., 2011; Dalkmann et al., 2012; Jechalke et al., 2015). Thus, how and which secondary anthropogenic activities affect the prevalence of ARGs in subsoil and groundwater environments remains a major gap of knowledge. Böckelmann et al. (2009) reported the presence of tetracycline and erythromycin ARGs in the groundwater of MAR sites, without any clear trend or impact associated with MAR. In the groundwater of a MAR site in Israel, *bla*_TEM_ and *qnrS* were below the limit of quantification (LOQ) (Elkayam et al., 2017). However, Lüneberg et al. (2017) demonstrated that during TWW irrigation, the prevalence of ARGs increased in the water flow paths towards the aquifer of soil/subsoil microcosms spiked with antibiotics. Taking into account that subsoil and groundwater microbial communities from various locations exhibit high variation and complexity (Haack et al., 2004; Kumar et al., 2017; Anantharaman et al., 2016; Yan et al. 2020), the number of previous studies is insufficient to cover the gap of knowledge regarding the influence of TWW irrigation on ARG prevalence in subsoil, subsoil pore-water and groundwater microbial communities.

Thus, the objective of the present study was to systematically investigate the impact of TWW irrigation on ARG prevalence in subsoil pore-water. Our working hypothesis was that TWW irrigation increases the prevalence of ARGs in subsoil pore-water. To test this hypothesis, we sampled the percolated subsoil pore-water in lysimeter-wells of a real-scale agricultural field, regularly subjected to TWW irrigation for commercial farming of crops. Samples were taken regularly across a one-year sampling period, across different periods and intensities of irrigation and after a long irrigation break, to gain insights into how TWW irrigation affects ARG prevalence on the field scale. Samples were taken at three different depths to ensure that observed TWW irrigation effects are consistent across the full soil depths profile and not restricted mainly to upper soil layers. All samples were analysed with qPCR for the abundance of six ARGs, the gene *intI1*, which is associated with mobile genetic elements (MGE) and an indicator for anthropogenic pollution (Gillings et al., 2015), and the gene 16S rRNA.

The interpretation of long-term, real-scale, field-based observations is however regularly affected by the variability of diverse environmental parameters, such as temperature or precipitation. To overcome this limitation, we here combined our field study with lab-based mesocosm experiments under controlled conditions. We set up subsoil pore-water mesocosms irrigated with either freshwater or TWW, to comprehensively test the validity of the insights gained from the real-scale field study, hence elucidating the impact of TWW irrigation on subsoil pore-water ARG prevalence.

## 2. Materials and methods

### 2.1 Sampling campaign of the lysimeter-wells

The sampled sandy (cambisol), agricultural field is located in Wendeburg, Germany (N: 52.359500, E: 10.399833; Fig. S1) and is associated with the Braunschweig Wastewater Association (BWA). Further characterization of the field site (e.g. soil composition) has been provided through previous studies (Ternes et al., 2007).

The specific field contained pre-installed lysimeter-wells that allowed the collection of percolated water from three different depths: 40, 80 and 120 cm (Fig. S2). Every year a different schedule of TWW irrigation with various intensities of irrigation is applied to the field, according to agricultural plans of the farmers and amount of natural precipitation. The field is irrigated with TWW or TWW mixed with digested sludge (TWW & DS), collected from the Urban Wastewater Treatment Plant (UWTP) of BWA, depending on the nutrient-demand of the grown crops. We sampled soil pore-water from the lysimeter-wells during several periods of irrigation intensity for eleven months: high-intensity irrigation, intermediate-intensity irrigation, low-intensity irrigation and after a long irrigation break (Table 1).

**Table 1:** Sampling dates, conditions and irrigation intensity of the lysimeter-wells (TWW: Treated Wastewater, DS: Digested Sludge).

| Sampling | Irrigation type | Conditions | Irrigation intensity |
| --- | --- | --- | --- |
| Apr-17 | TWW & DS | Continuous and regular irrigation was initiated on the field. | High |
| May-17 | TWW & DS | The field was irrigated continuously and regularly and was already irrigated for one month. | High |
| Jun-17 | TWW & DS | The field was irrigated continuously and was already irrigated for two months. | High |
| Oct-17 | TWW | The field was irrigated spontaneously. | Intermediate |
| Feb-18 | No-Irrigation<br>(Rainfall) | The field was not irrigated since October 2017 and was only exposed to natural precipitation. | Irrigation break |
| Feb-18 | TWW | The field was irrigated for the first time after the irrigation break. | Low |
| Mar-18 | TWW | The field was irrigated regularly from February. | Low |

Samples of percolated water were collected in sterilised 2 L glass bottles at each lysimeter-well’s percolation-point (Fig. S2). The bottles were placed in the percolation-points prior to an irrigation event and percolated water was collected for three days after each event (4 samples per depth). At the end of the irrigation break (October 2017-February 2018), the bottles were placed on the lysimeters to equally collect percolated water from natural soil-moisture. Samples were stored on ice and transferred to the lab. Bacteria were captured by filtration of 0.1 to 0.5 L of percolated water (depending on collected volume) in triplicate per sample within 48 hrs. The filters (polycarbonate, 0.2 μm pore size, 47 mm diameter, Sartorius, Germany) were stored at −20°C prior to DNA extraction.

### 2.2 Subsoil pore-water mesocosms

For the mesocosm experiments, we sampled forest soil, located 10-20 m adjacent to the TWW irrigated field (Table S2 describes the physicochemical characteristics of the forest soil). The samples were taken from 12 individual sampling points at 0-60 cm depth. After air-drying, and sieving (~6 mm mesh size), the soil was homogenised to create a composite soil sample that was used for the mesocosm experiments. Moreover, soil samples were preserved for analysis of background levels of ARGs (n=12).

Mesocosms (Fig. S3) were assembled from cylindrical plastic tubes of 66 cm height and 4.5 cm radius. The bottom of each mesocosm (5 cm height) was filled with quartz gravels (size: ~3 mm^3^) and an inner tube (1.5 cm radius) was placed centrally to allow the collection of the subsoil pore-water (Fig. S3). Mesocosms were then filled with the previously homogenized soil up to 50 cm height, resulting in a total soil volume per mesocosm of 2,827.6 mL.

Mesocosms were divided into two groups, with four replicate mesocosms per treatment: The Freshwater-Group and TWW-Group were defined with respect to the type of irrigation water. TWW for irrigation was obtained from an UWTP nearby the Dresden area (Kaditz, Germany; N: 51.070640, E: 13.680888). The freshwater was collected from a shallow well (depth ~7 m) located next to the Elbe river, in Pirna, Germany (N: 50.965905, E: 13.924034). The mesocosms were irrigated with 350 mL of water, which led to saturation. Removal of residual water and renewed irrigation were performed aseptically three times per week.

Both groups were initially irrigated with freshwater for two weeks, to stabilise and equilibrate the soil conditions. Then the TWW-Group switched to TWW irrigation for three weeks, while the Freshwater-Group was continuously irrigated with freshwater. Subsoil pore-water samples (200 mL per mesocosm) were taken aseptically from each mesocosm of both groups. First sampling took place at the end of the two-week freshwater irrigation/stabilisation-period (Week 0), the second in the 1^st^ week after switching to TWW irrigation (Week 1) and the third in the 3^rd^ week after switching to TWW irrigation (Week 3). Bacteria were harvested from the samples through filtration as described above. Filters were frozen directly after sampling and stored at −20°C prior to DNA extraction. TWW and freshwater samples (0.5 L, n=6 per irrigation type) were equally filtered and stored for DNA extraction.

### 2.3 DNA extraction and quantitative real time PCR

We used the DNeasy PowerWater Kit and the DNeasy PowerSoil Kit (Qiagen, Germany) according to the manufacturer’s instructions to extract DNA from the water and soil samples, respectively. The quantity and quality of DNA was measured with NanoDrop (Thermo Fischer Scientific, Germany). The samples were analysed with quantitative real-time PCR (qPCR) for six ARGs (*sul1, bla*_OXA-58_*, tetM*, *qnrS*, *bla*_TEM_ and *bla*_CTX-M-32_), *intI1* and 16S rRNA. Reactions were performed in a MasterCycler RealPlex (Eppendorf, Germany) at final volume 20 μL with 10 μL of Luna Universal qPCR Master Mix (New England Biolabs, Germany). The concentration of the primers varied from 0.2-0.5 μM (for further details regarding genes, reagents, primers and temperature for each gene see Table S3). Standard curves were created during every qPCR run, with the use of the same plasmid vector and procedures as described previously (Cacace et al., 2019).

Standard curves with amplification efficiency 0.9-1.1 and R^2^ ≥ 0.99 were accepted and melting curve analysis was performed to assess the amplicons’ specificity. Screening for PCR inhibition was performed by spiking a plasmid containing a gene present in low abundance in our samples (*bla*_CTX-M-32_, spiking concentration 4*10^6^ copies/μL): no inhibition was detected in any of the samples. The absolute abundance of genes was finally expressed as gene copies/L and the relative abundance as the ratio of gene copies per copy of 16S rRNA.

### 2.4 Data processing and statistical analysis

Prior to any statistical analysis, every gene in every sample that was below LOQ was set as 1 copy/L for absolute abundance and 10^−8^ for relative abundance (one order of magnitude lower than the minimum relative abundance observed for any gene in any sample, ~10^−7^). Data was then log10-transformed before performing statistical analysis. The programing language R (www.R-project.org, v. 3.5.3) was used for graphical representations, with the packages “ggplot” (Whickham, 2016) and “ggpubr” (v. 0.2.2, Kassambara, 2019). Significant differences were assessed with the Wilcoxon rank sum test or in case of group comparisons with the Kruskal-Wallis test (package “ggpubr”). A post-hoc test (Dunn’s test with Benjamini-Hochberg correction) was performed with the use of the package “dunn’s test” (v1.3.5, Dinno, 2016), to assign significant differences from pairwise multiple comparisons. The difference in ARG profiles of the lysimeter-well samples were assessed with PERMANOVA tests (“adonis” function, method=“euclidean”) from the “vegan” package (v2.5-6, Oksanen et al., 2011). In addition, we performed pairwise comparisons for the field and the mesocosm groups over time with the “pairwiseAdonis” package (Martinez Arbizu, 2019; v0.3, function “pairwise.adonis”, method=“euclidean”) along with Benjamini-Hochberg correction for multiple comparisons.

## 3. Results

### 3.1 Lysimeter-Wells

#### 3.1.1 Seasonal variation, rather than TWW irrigation affects subsoil pore-water bacterial abundance

Throughout the sampling period, the studied field site was irrigated with either TWW or TWW mixed with digested sludge (DS). The bacterial load in the irrigation water, determined through 16S rRNA abundance, remained stable, whether there was a mixing with DS or not: 9.8±0.36 log10 copies/L for TWW & DS (April, May, June 2017) and 9.9±0.34 log10 copies/L for the TWW (October 2017, February and March 2018) (Fig. 1) (Wilcoxon rank sum test, p=0.16, n=3).

**Figure 1:**
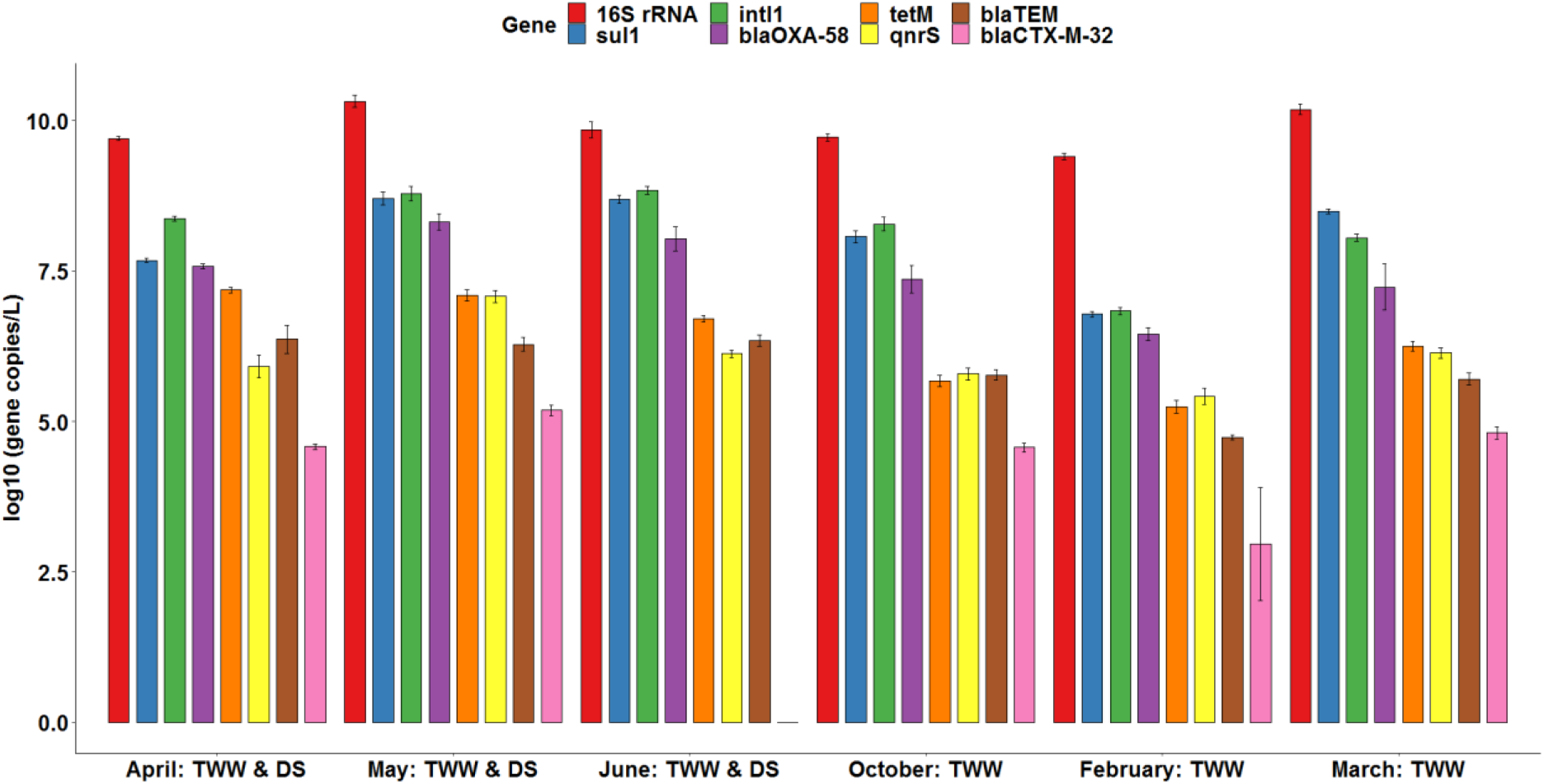
Absolute abundance (copies/L) of the selected genes in the irrigation water of BWA. TWW: Treated Wastewater, DS: Digested Sludge (n=3).

The absolute abundance of 16S rRNA gene copies in the pore-water samples throughout the year ranged from 8.0-9.8 log10 copies/L (Fig. 2), while we observed similar patterns of in-/decrease at all three different depths. During intensive irrigation periods (April, May and June 2017), the 16S absolute abundance was constantly one order of magnitude lower at each sampled depth compared to the rest of the sampling campaign (Fig. 2). For example at 120 cm depth, the abundance of 16S rRNA was 7.9-8.3 log10 copies/L in April-June 2017 (periods of high-intensity irrigation). Yet, in October 2017, Feb. (1) 2018 (before start of TWW irrigation) and Feb. (2) 2018 (after the restart of TWW irrigation) the abundance of 16S rRNA was significantly higher at 9.0±0.2, 8.9±0.1 and 8.9±0.3 log10 copies/L, respectively (Fig. 2, p= 2.2 × 10^−14^, Kruskal-Wallis, n=4).

**Figure 2:**
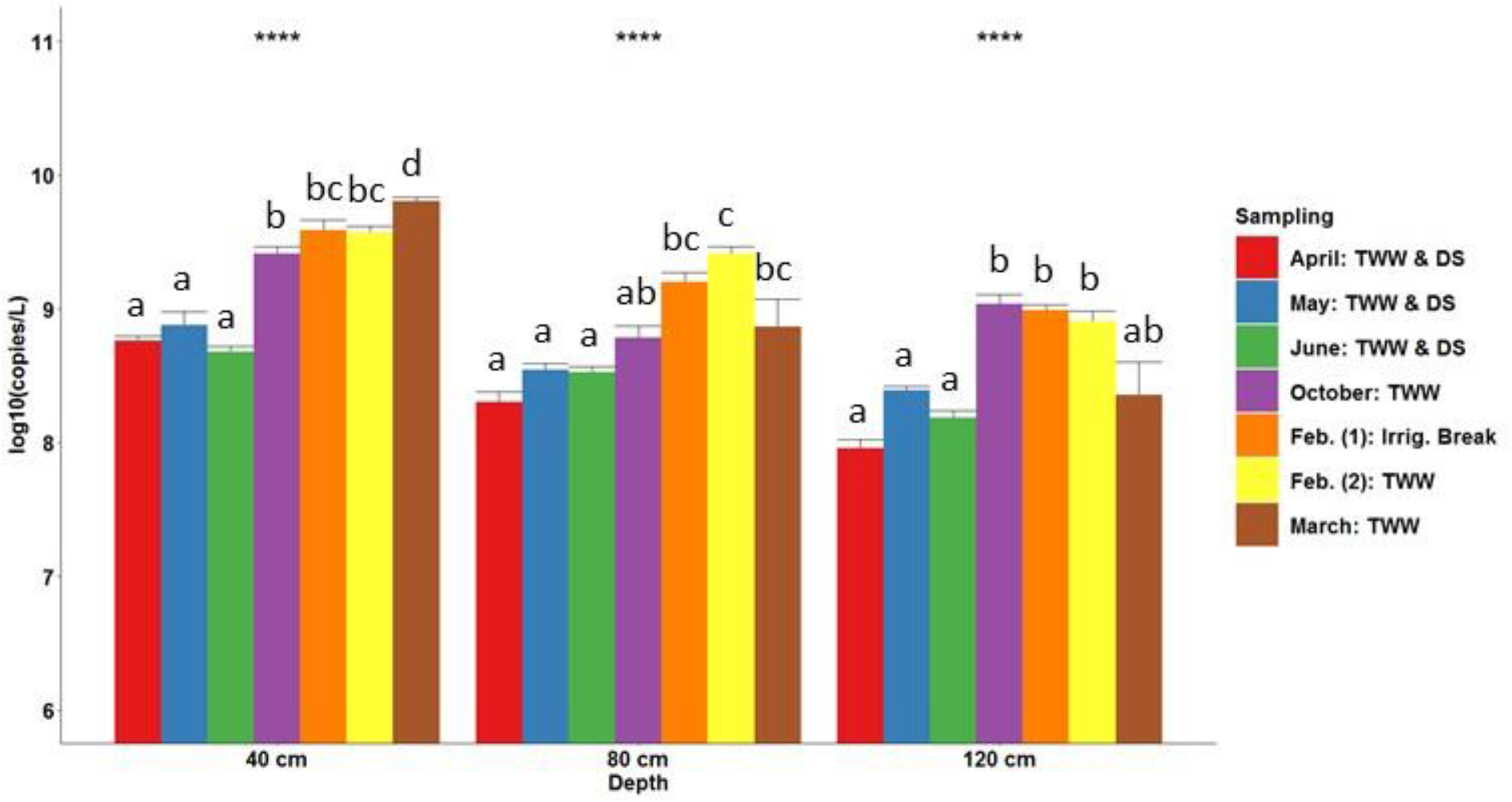
Absolute abundance (log10 copies/L) of 16S rRNA in percolated subsoil pore-wate r from the lysimeter-wells during periods of high (April, May, June), moderate (October) and low intensity irrigation (Feb. (2), March) or irrigation break (Feb. (1)). Kruskal-Wallis test: *p<0.05, p<0.01, *p< 0.001, p<0.0001, n=4. The letters “a” to “d” were assigned to non-signific antly different groups after pairwise comparisons with Dunn’s test along with Benjamini-Hoc hberg correction and cutoff p<0.05. TWW: Treated TWW, DS: Digested Sludge, Irrig.: Irriga tion.

#### 3.1.2 Relative abundance of *sul1* and *intI1* increase with TWW irrigation intensity

All investigated ARGs as well as the *intI1* gene were detected in the irrigation TWW (Fig. 1). The genes with highest abundance in the irrigation water were *sul1* and *intI1.* Their mean relative abundance was −1.6 and −1.8 log10 copies/16S rRNA, respectively, while that of the any of the other ARGs varied from −3 to −7 log10 copies/16S rRNA. The rank abundance in the irrigation water was determined as *intI1* > *sul1* > *bla*_OXA-58_ > *tetM* > *qnrS* > *bla*_TEM_ > *bla*_CTX-M-32_ (Fig. 1).

All investigated genes were also present in subsoil pore-water, with *sul1* and *intI1* again being the most abundant (*intI1:* −2.7±1.0 log 10 copies/16S rRNA, *sul1*: −2.9±0.5 log 10 copies/16S rRNA). To determine if irrigation intensity affects the ARG profile of the subsoil pore-water, we grouped the different samples based on the intensity of irrigation during the sampling period (high, intermediate, low, irrigation break). ARG profiles of every group of irrigation intensity were significantly different (p=0.001 with R^2^<0.25, Euclidean Distance, Benjamini-Hochberg correction) (Table 2), based on pairwise multiple comparison with PERMANOVA tests, based on the relative abundance of ARGs and *intI1*.

**Table 2:** Pairwise comparisons of the lysimeters groups over periods of irrigation with PERMANOVA test (pairwise.adonis function, Euclidean distance), with Benjamini-Hochberg correction for multiple comparisons.

| Pairwise Comparisons | R<sup>2</sup> | Adjusted P-Value |
| --- | --- | --- |
| High Intensity Period vs Intermediate Intensity Period | 0.060169 | 0.001 |
| High Intensity Period vs Irrigation Break | 0.214535 | 0.001 |
| High Intensity Period vs Low Intensity Period | 0.168915 | 0.001 |
| Intermediate Intensity Period vs Irrigation Break | 0.214919 | 0.001 |
| Intermediate Intensity Period vs Low Intensity Period | 0.075231 | 0.001 |
| Irrigation Break vs Low Intensity Period | 0.033387 | 0.001 |

To gain insights into which genes most significantly influenced the respective differences in ARG and *intI1* profiles, we compared the relative abundance of each gene, in relation to irrigation intensity. Trends observed for ARGs and *intI1* were unless otherwise stated consistent across the three different depths sampled (e.g. Fig. 3), hence examples given for observed patterns below will be restricted to profiles obtained at a single depth. Specifically, the relative abundance of the two most abundant genes, *sul1* and *intI1,* exhibited a positive correlation with the intensity of TWW irrigation (Fig. 3). In general, the relative abundance of both genes at each of the sampled depths was higher during the period of intensive irrigation (April-June 2017), compared to periods of less irrigation (Dunn’s test, p<0.05, n=4). For example, the relative abundance of *sul1* at 120 cm depth significantly differed among the various periods of irrigation (p= 2*10^−16^, Kruskal-Wallis test, n=4). It remained relatively stable from April to June 2017, where continuous irrigation with TWW & DS was performed, from −2.6±0.1 to −2.3±0.5 log 10 copies/16S rRNA. In October 2017, after the first irrigation break took place, the relative abundance of *sul1* significantly decreased to −3.1±0.3 log10 copies/16S rRNA and further to −3.7±0.4 log10 copies/16S rRNA in February 2018 (end of 2^nd^ irrigation break), one order of magnitude lower than April-June 2017. Similar seasonal trends, in relation to irrigation intensity, were observed for *sul1* and *intI1* at all depths. The only difference in trends was detected immediately upon restarting TWW irrigation in February 2018. Relative abundances for both genes at all depths remained around one order of magnitude lower compared to the intense irrigation period in June 2017. However, a clear trend of increasing relative abundance of *sul1* and *intI1* was detected after irrigation restarted except for *intI1* at 120 cm depth, where no immediate impact of restarting irrigation.

**Figure 3:**
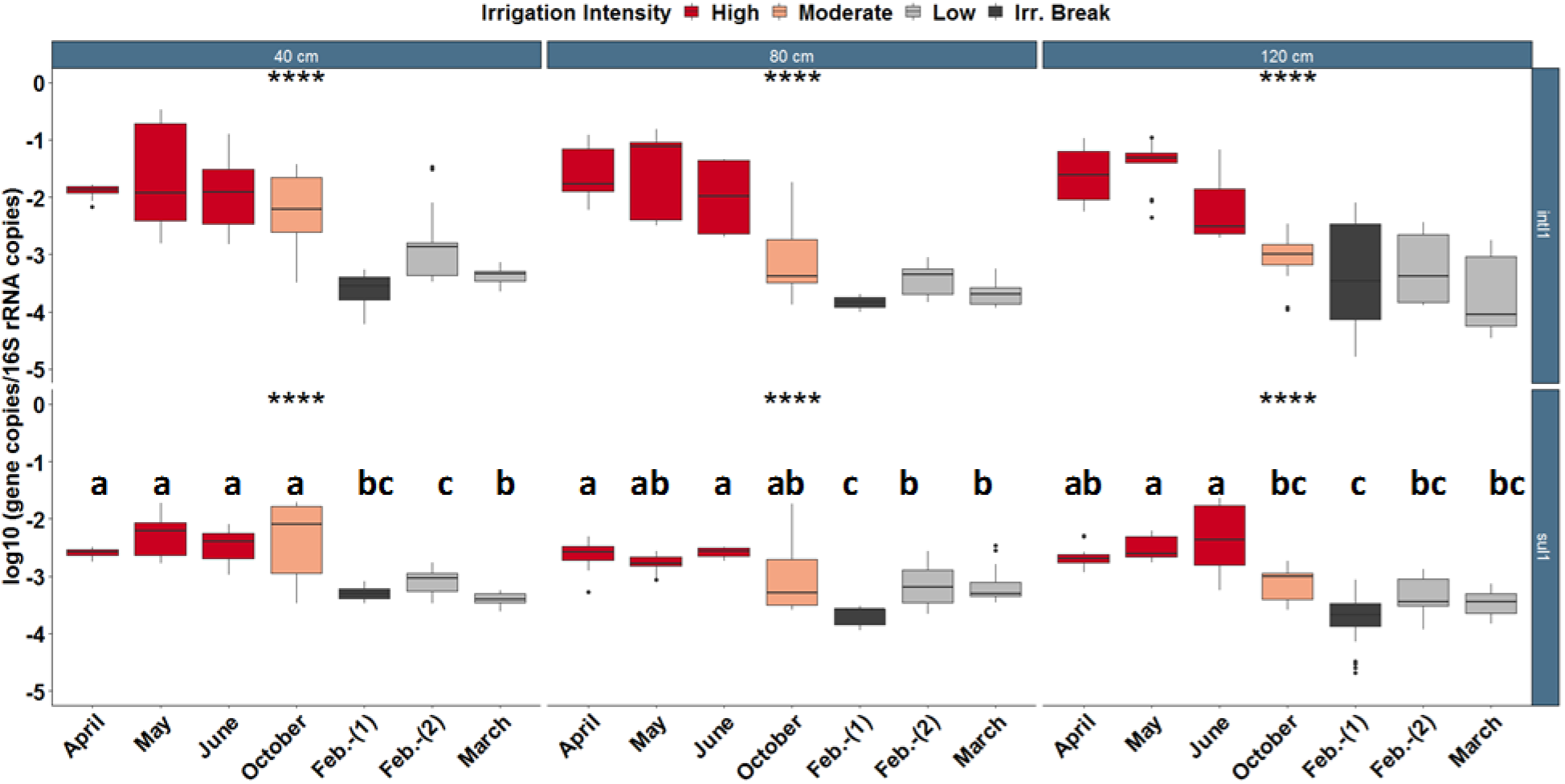
Relative abundance of *sul1* and *intI1* during periods of high, moderate, low irrigation intensity or irrigation break (Irr. Break), assigned based on the level of irrigation from Table 1. Kruskal-Wallis test: *p<0.05, **p<0.01, p< 0.001, *p<0.0001, n=4. The letters “a” to “c” were assigned to non-significantly different groups after pairwise comparisons with Dunn’s test along with Benjamini-Hochberg correction and cutoff of p<0.05.

Among the other genes, *qnrS* and *bla*_OXA-58_ exhibited similar increases in relative abundance after restarting TWW irrigation in February 2018 (Fig. S4 and S5): e.g. *qnrS* (Fig. S4) at 120 cm depth increased from −6.4±0.4 to −5.7±0.3 log10 copies/16S rRNA (p=0.02, Wilcoxon rank sum test, n=4). Despite this increase, they did not display the stable trend observed for *sul1* and *intI1,* where higher relative abundance is correlated with periods of intensive TWW irrigation (Fig. S4 & S5). None of the other genes (*bla*_CTX-M-32_, *bla*_TEM_, *tetM*) exhibited any clear trends in relation to irrigation intensity (Fig. S6, S7 & S8). For example, despite continuous intense irrigation with TWW & DS between April and June 2017, the relative abundance of *bla*_TEM_ was two orders of magnitude lower in June 2017 compared to April 2017.

While the sampling of lysimeter-wells provided clear indications that TWW irrigation seemed to affect the prevalence of ARGs in the subsoil pore-water, there were high variations of physical conditions (e.g. temperature, precipitation) in the investigated field across the year. Such variation is common among field studies, thus insights gained from field studies should to be tested further under controlled conditions.

### 3.2 Mesocosm experiments

#### 3.2.1 Absolute bacterial abundance in subsoil pore-water is independent of the irrigation water and its bacterial load

By comparing the impact of TWW or freshwater irrigation, the mesocosm experiments aimed at verifying the effects of TWW irrigation on ARG abundance subsoil pore-water, obtained from the field study under controlled conditions. The absolute bacterial abundance significantly differed between the two types of irrigation water (9.7±0.2 log10 16S rRNA copies/L TWW; 7.2±1 log10 copies/L Freshwater; n=6; Wilcoxon rank sum test, p=3.6 × 10^−5^, n=6) (Fig. S10A, Fig. 4A). The initial 16S rRNA abundance in the pore water of the mesocosms after 3 weeks of equilibration with freshwater irrigation was 10.6±0.2 log10 copies/L for the Freshwater-Group and 10.4±0.2 log10 copies/L for the TWW-Group in Week 0, approximately three orders of magnitude higher than in the freshwater irrigation feed (Fig. 4A). No significant difference between the two groups in absolute 16S abundance was observed after irrigation switched to TWW in half of the mesocosms. In both groups 16S abundance decreased to 9.8±0.2 and 9.7±0.3 log10 copies/L in Week 1 (Dunn’s test, p<0.05, n=4), yet no significant difference between the two groups of mesocosms was detected (Dunn’s test, p>0.05, n=4). However, the Freshwater-Group displayed slightly higher 16S rRNA abundance than the TWW-Group in Week 3 (Freshwater: 10±0.1 log10 copies/L; TWW: 9.7±0.2 copies/L, Dunn’s test, p<0.05, n=4). Therefore, continuous irrigation with higher bacterial loads through TWW application did not increase the absolute abundance of 16S rRNA in the pore water of the TWW-Group. On the contrary, 16S rRNA abundance showed peculiar and rather stochastic dynamics, since there was an almost ten-fold decrease from Week 0 to Week 1 (one-week difference) for both types of irrigation (Fig. 4A, Dunn’s test, p<0.05, n=4).

**Figure 4:**
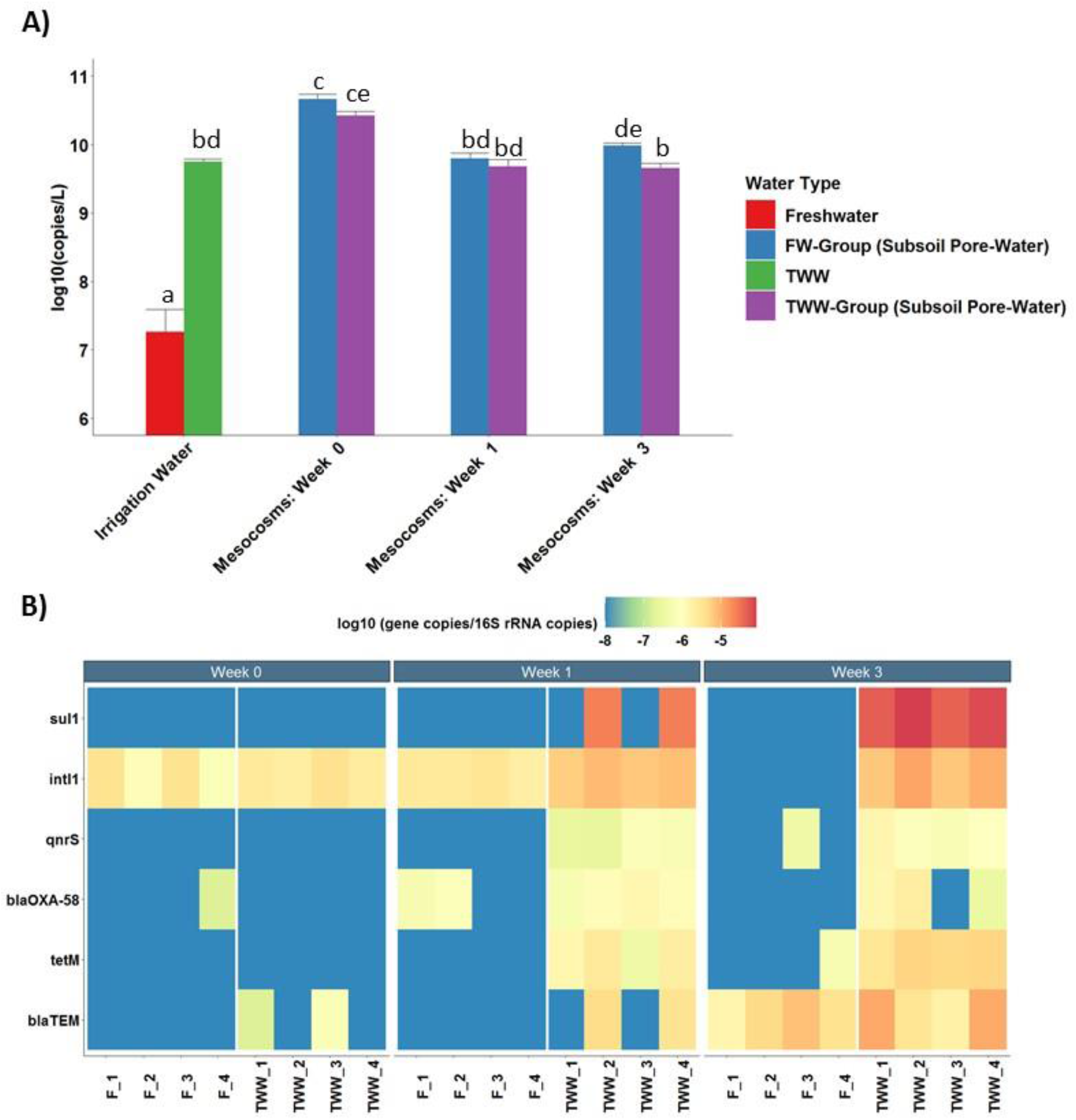
A) Absolute abundance (log10 gene copies/L) of 16S rRNA in the percolated subsoil pore-water of the mesocosms and the respective irrigation waters. Irrigation water: n=6; Mesocosms subsoil pore water: n=4. The letters “a” to “d” were assigned to non-significantly dif ferent groups after pairwise comparisons with Dunn’s test along with Benjamini-Hochberg correction and cutoff p<0.05. TWW: Treated Wastewater, FW: Freshwater. B) Relative abundance of ARGs and *intI1* (log10 gene copies/16S rRNA) in the subsoil pore-water of mesocosms (F: Freshwater irrigated, TWW: Treated Waste Water irrigated). The gen e *bla*_CTX-M-32_ was not shown as it was below LOQ in all samples. Pairwise comparisons with P ERMANOVA test, along with Benjamini-Hochberg correction, can be found in Table 3.

#### 3.2.2 TWW irrigation increases the relative abundance of *sul1, intI1, qnrS, tetM* and *bla* OXA-58 in subsoil pore-water of mesocosms

Upon initiation of the mesocosms experiments, the ARG-profile of the soil was analysed. While, *sul1, tetM, qnrS* and *bla*_OXA-58_ were below the LOQ, *bla*_TEM_, *bla*_CTX-M-32_ and *intI1* were present in the soil (Fig. S9). In addition, the ARG-profile of both selected irrigation water sources (freshwater and TWW) was analysed. All analysed genes were present in both, TWW and freshwater but constantly exhibited higher absolute abundances in TWW than freshwater (Wilcoxon rank sum test, p<0.0001, n=6, Fig. S10A). The relative abundance of ARGs was higher in TWW as well (Wilcoxon rank sum test, p<0.0001, n=6), with the exception of *bla*_CTX-M-32_ and *bla*_TEM_, where no significant difference between the two irrigation types was observed (Wilcoxon rank sum test, p>0.05, n=6, Fig. S10B).

**Table 3:** Pairwise comparisons of the ARG and *inti1* profiles in the mesocosm percolated wat er with PERMANOVA test along with Benjamini-Hochberg correction for multiple comparis ons (FW: Freshwater irrigated, TWW: Treated Wastewater irrigated, W: Week).

| Pairwise Comparisons | R<sup>2</sup> | Adjusted p-value |
| --- | --- | --- |
| FW-W0 vs TWW-W0 | 0.106597 | 0.285 |
| FW-W0 vs FW-W1 | 0.053469 | 0.285 |
| FW-W0 vs TWW-W1 | 0.447706 | 0.005 |
| FW-W0 vs FW-W3 | 0.715021 | 0.005 |
| FW-W0 vs TWW-W3 | 0.689762 | 0.005 |
| TWW-W0 vs FW-W1 | 0.133986 | 0.285 |
| TWW-W0 vs TWW-W1 | 0.425895 | 0.005 |
| TWW-W0 vs FW-W3 | 0.649432 | 0.005 |
| TWW-W0 vs TWW-W3 | 0.66361 | 0.005 |
| FW-W1 vs TWW-W1 | 0.416059 | 0.005 |
| FW-W1 vs FW-W3 | 0.688689 | 0.005 |
| FW-W1 vs TWW-W3 | 0.666517 | 0.005 |
| TWW-W1 vs FW-W3 | 0.522884 | 0.005 |
| TWW-W1 vs TWW-W3 | 0.186126 | 0.132 |
| FW-W3 vs TWW-W3 | 0.66171 | 0.005 |

After both groups of mesocosms were initially equilibrated through freshwater irrigation, the profile of ARGs and *intI1* was identical in Week 0 (Fig. 4B, Table 3, PERMANOVA, p>0.05, n=4). However, after the switching half the mesocosms to TWW irrigation, the ARG-profile of the two groups differed significantly in Week 1 (PERMANOVA, p=0.005, R^2^=0.42) and Week 3 (PERMANOVA, p=0.005, R^2^=0.66) (Fig. 4B, Table 3). For example, the relative abundance of *intI1* of the TWW-Group in Week 0 was −5.6±0.3 log10 copies/16S rRNA and in Week 1 significantly increased to −5.2±0.2 log10 copies/16S rRNA (Kruskal-Wallis test, p=0.003, n=4) after which it remained stable in Week 3 at −5.1±0.2 log10 copies/16S rRNA. In the Freshwater-Group, the relative abundance of *intI1* decreased from −5.7±0.3 log10 copies/16S rRNA in Week 0 to below LOQ after Week 3 (Kruskal-Wallis test, p=0.0003, n=4). Thus, continuous freshwater irrigation led to a significant reduction up to elimination of *intI1* from the subsoil pore-water. Furthermore, *sul1* was not detected in Week 0 in either group or at any time during freshwater irrigation. However, *sul1* was detected in two of the four mesocosms of the TWW-Group in Week 1 and finally in all mesocosms of the TWW-Group in Week 3 (−4.2±0.3 log10 copies/16S rRNA), resulting in a significant difference of relative abundance compared to the Freshwater-Group (p=0.00041, Wilcoxon rank sum test, n=4). Similarly, *tetM* was mainly detected in the TWW-Group following similar trends as *intI1* and *sul1* during the weeks of irrigation.

The gene *bla*_CTX-M-32_ was not detected, in either group in all weeks of irrigation. In addition *bla*_TEM_ was below LOQ for the majority of samples in Week 0, but its relative abundance increased to 6.5 log10 copies/16S rRNA in both groups of irrigation in Week 3. Therefore, TWW irrigation did not influence the dissemination of *bla*_TEM_ in the subsoil pore-water relative to freshwater irrigation. The genes *qnrS* and *bla*_OXA-58_ were also below LOQ in Week 0 (both groups), but were detected in Week 1 exclusively in the TWW-Group. However, their relative abundance remained stable once detected from Week 1 to Week 3. For example, relative abundance of *qnrS* was present at −6.6±0.7 log10 copies/16S rRNA copies/16S rRNA at Week 1 and −6.1±1 log10 copies/16S rRNA at Week 3 (p=0.42, Wilcoxon rank sum test, n=4, Fig. 4B). In contrast, *qnrS* and *bla*_OXA-58_ remained below LOQ for the majority of samples from the Freshwater-Group throughout the experiment. Therefore, TWW irrigation showed a strong influence on the dissemination of *sul1* and *intI1* and a moderate influence for *qnrS* and *bla*_OXA-58_, which was in line with the results from the lysimeter-wells. The more controlled and consequently more sensitive mesocosm experiments were able to further document effects of TWW irrigation for *tetM,* when compared with freshwater irrigation

## 4. Discussion

In this field study, the prevalence of ARGs and *intI1* in the subsoil pore-water of a real-scale agricultural field, subjected to TWW irrigation for commercial farming operation, was positively correlated with TWW irrigation intensity for the genes *sul1*, *intI1, qnrS* and *bla*_OXA-58_. This trend and hence a causal link was later confirmed in controlled lab-scale mesocosm experiments, overcoming the lack of representative controls (freshwater irrigated fields with lysimeter-wells) and varying environmental conditions (temperature, precipitation). Therefore, our working hypothesis that TWW irrigation promotes the spread of ARGs and *intI1* into subsoil pore-water was confirmed. To the best of our knowledge, this is the first study that investigated the impact of TWW irrigation on the profile of ARGs in this so far underappreciated matrix through the combination of real-scale TWW-irrigated field investigations and mesocosm experiments.

The impact of TWW irrigation on the relative abundance of ARGs and *intI1* displayed consistent but not completely identical patterns at the three different sampled depths of our real scale field study. All depths were equally affected during high intensity irrigation, with similarly increased relative abundance of *sul1* and *intI1* at all three depths during high intensity irrigation periods. Therefore the effect of TWW irrigation is not only restricted to the subsoil pore-water of upper, but also carried through to the deeper soil layers. The main exception we observed was *intI1* only increasing in the two upper soil layers immediately upon the restart of low intensity irrigation after a prolonged irrigation break. This indicates a temporal delay in exposure to and hence effect of TWW irrigation in the deeper soil layers.

The fact that not all ARGs and *intI1* followed parallel trends of in- and decrease correlated with irrigation intensity, leads to the assumption that the abundance of ARGs in soil pore-water is not exclusively due the leaching of irrigation water, a process described previously as a main source of the microbial load in lysimeter samples (Forslund et al., 2011). Assuming such a leaching scenario would rather lead to a uniform, parallel increase of all ARGs, based on their relative abundance in irrigation water as a function of irrigation intensity. Hence, a complex combination of interactions of soil, TWW and their respective microbiomes needs to be taken into account when evaluating the effect of TWW irrigation on ARG abundance in soil pore-water. These include microbial ecological and evolutionary processes such as horizontal gene transfer of resistance genes, competition and selection dynamics, but also physico-chemical processes like transport and leaching.

A strong indicator for horizontal gene transfer between TWW and soil microbes playing a major role in the observed increase in resistance genes in the subsoil pore-water is that among the genes tested, *sul1* and *intI1* showed the highest relative abundance and strongest correlation with irrigation intensity. The integrase gene *intI1,* commonly used as an indicator for anthropogenic pollution (Gillings et al., 2015), is generally associated with MGE and usually located in close genetic proximity to ARGs or other genes connected to adaptive stress responses in plasmids or transposons (Gillings, 2017). For example, *sul1* has been frequently found as part of *intI1* gene cassettes that are located on mobile genetic elements and conjugative plasmids (Gillings et al., 2015; Gillings et al., 2018), which have the ability to transfer to a the majority of the diverse bacterial phyla found in soil (Klümper et al. 2015). Consequently, their parallel increase in abundance is most likely connected to them being co-located on MGEs that get co-transferred and co-selected. Unsurprisingly, these mobile genes were also highly abundant in TWW effluents from several geographical regions (Cacace et al., 2019; Pärnänen et al., 2019).

While these genes were also present in the freshwater used for irrigation of mesocosms, their abundance was orders of magnitude lower, compared to TWW. However, taking into account the high horizontal mobility of *sul1* and *intI1* (Gillings et al., 2015; Gillings et al., 2018) we presume that long-term freshwater irrigation could in theory lead to an enrichment of *sul1* in the soil and hence the subsoil pore-water. In a previous study *sul1* was one of the ubiquitously detected ARGs in six groundwater wells (Szekeres et al. 2018), suggesting that its dissemination in the subsoil/groundwater microbiome is indeed rather successful. However, *intI1*, which was initially detected in the soil after the two weeks of equilibration with freshwater (Week 0), was subsequently eliminated from mesocosms in the Freshwater-Group after five weeks of total freshwater irrigation (Week 3), indicating that in addition to its gene transfer potential, positive selection might be needed for enrichment in the subsoil pore-water.

The presence of selective agents may cause such positive selection for ARGs and/or promote horizontal gene transfer of MGEs by inducing the bacterial stress response. These agents can for example include pharmaceutical residues (including antibiotic and non-antibiotic residues) (Lin & Gan, 2011; Dalkmann et al., 2012; Manaia et al., 2018b) or heavy metal ions (Klümper et al., 2017; Zhang et al., 2018; Zhao et al., 2019). The gene *sul1* confers resistance to sulphonamides, a class of antibiotics of synthetic origin, which have been reported to occur in high concentration in TWW (Nikolaou et al., 2007; Barnes et al., 2008; Avisar et al., 2009; Michael et al., 2013) and they persist in sub-surface environments (Barnes et al., 2008; Avisar et al., 2009). For example, sulphamethoxazole was detected in groundwater from different areas in the USA (Barnes et al., 2008) and in phreatic aquifer samples in Israel (Avisar et al., 2009). In addition, the reported maximum values of sulphamethoxazole concentrations reported for these groundwater samples was 1.1 μg/L (Barnes et al., 2008) thus exceeding the predicted no effect concentration for the selection of sulphamethoxazole resistance for environmental strains (0.5 μg/L), based on predictive models (Bengtsson-Palme & Larsson, 2016). Lüneberg et al. (2017) showed that during limited and simulated TWW irrigation events, the prevalence of *sul1* indeed increased in the water flow paths towards the aquifer of their tested soil/subsoil mesocosms as a function of sulphamethoxazole concentrations.

Further, non-antibiotic pharmaceuticals of TWW origin, such as carbamazepine, can induce the mobilisation of ARGs and MGE (Wang et al., 2019). Carbamazepine is one of the most common non-antibiotic drugs, regularly found in TWW (Clara et al., 2004; Nikolaou et al., 2007; Gasser et al., 2011), while it has shown high persistence in groundwater as well (Clara et al., 2004; Gasser et al., 2011; Sui et al., 2015). The preferential increase of specific mobile ARGs and *intI1* observed in our study could probably be explained by the fact that these agents may be released by TWW irrigation and are able to persist in the subsurface environments as well. Ternes et al. (2007) reported the presence of pharmaceutical residues including sulphamethoxazole (maximum value: 0.12 μg/L) and carbamazepine (maximum value: 1.3 μg/L) in the exact lysimeter-wells of the same TWW irrigated field we studied. Based on our findings and previous results we presume that a causal link might exist between the dissemination/invasion success of *sul1,* the introduced dose of *sul1* and the combination of selective agents, which are originally co-introduced into the subsoil/groundwater matrices through TWW irrigation. Therefore, further controlled exploration of *sul1* dissemination in subsoil/groundwater, in combination with the presence and persistence of selective agents in these matrices, is necessary to elucidate the potential mechanisms behind this link.

The subsoil pore-water in the field and mesocosms showed high 16S rRNA absolute abundance, but was not influenced by the type or intensity of irrigation, indicating that physical processes such as direct leaching of the irrigation water into the subsoil pore-water plays only a minor role. In the field, we observed a seasonal rather than irrigation intensity based trend of 16S rRNA absolute abundance; the highest absolute abundance occurred during the irrigation break and low intensity irrigation. In addition, the bacterial load did not increase after the restart of irrigation in February. In the mesocosms, the subsoil pore-water exhibited 3-4 orders of magnitude higher 16S rRNA absolute abundance than the freshwater, and still 0.5-1 orders of magnitude higher abundance than the irrigation TWW. Despite the fact that the freshwater had 2-3 orders of magnitude lower 16S rRNA abundance compared to TWW, no increase of 16S rRNA abundance in the subsoil pore-water was observed after switching to TWW irrigation. Therefore, the higher bacterial load added through TWW irrigation did not affect the bacterial load in the subsoil pore-water. A significant proportion of bacteria that inhabit the soil can mobilise and occupy the water matrix as well, during the percolation of water (Dibbern et al., 2014; Hermann et al., 2019). These indigenous soil/subsoil bacteria are well adapted to their specific niches (Marano et al., 2019; Cerqueria et al., 2019; Bahram et al., 2018; Jansson & Hofmockel, 2018) and thus most likely outcompete the invading TWW bacteria. Hence, soil bacteria are expected to make up the majority of the bacterial load in the subsoil pore-water.

In the present study, we examined the impact of TWW irrigation on the prevalence of ARGs in subsoil pore-water, however, TWW irrigation is not the only source of antibiotics, ARB and ARGs in most agricultural fields. Specifically, biosolids (Mathews & Reinhold, 2013) and manure amendment (Luby et al., 2016; Yang et al., 2016; Barrios et al., 2020) introduce a high loads of antibiotics and ARGs (Chen et al., 2016; Jechalke et al, 2016; Zhou et al., 2017; McKinney et al., 2018) and therefore might mask the impact of wastewater irrigation in this specific matrix in many cases (Cerquiera et al., 2019c). Further, only relative effects on ARG abundance can be measured as ARGs naturally occur at low levels in many environments of low anthropogenic or even pristine nature (D’Costa et al., 2011; Martinez, 2012, Gatica et al., 2016). In fact, our freshwater, sampled from a groundwater well near an agricultural suburban area of Dresden, also contained diverse ARGs such as *sul1.* However, its relative abundance was orders of magnitude lower compared to that detected in TWW. The TWW that was used for irrigation, both in the field (BWA) as well as the mesocosms (TWW Kaditz) is only subjected to secondary treatment. Advanced wastewater treatment leading to the reduction of ARGs from the effluents might minimise the differences between the profile of ARGs of TWW and freshwater along with the impact of TWW that we observed in our study (Michael et al., 2013; Manaia et al., 2018).

Overall, TWW irrigation increased the relative abundance of specific genes associated with antimicrobial resistance, mainly *sul1* and *intI1.* Combining mesocosm approaches with long-term studies on a regularly real-scale, commercially operated TWW irrigated field, proved a successful research tool to study irrigation effects in this so-far understudied environment. Still, further research into the mechanisms of the observed dissemination of ARGs in subsoil pore-water is necessary to improve the understanding of the ecology of ARGs in relation to freshwater/TWW irrigation, or other disturbances such as soil amendment in this matrix. This will allow generating mitigation strategies to minimize the risk associated with ARG dissemination in the subsoil pore-water and potentially in the groundwater during TWW irrigation scenarios.

## 5. Abbreviations

ARB: Antibiotic Resistant Bacteria
ARGs: Antibiotic Resistant Genes
As: Antibiotic Residues
BWA: Braunschweig Wastewater Association
DS: Digested Sludge
MGE: Mobile Genetic Elements
TWW: Treated Wastewater
UWTP: Urban Wastewater Treatment Plant

## 6. Acknowledgements

The authors would like to thank the executive director of AVBS, Bernhard Teiser for his assistance in providing access in the field and the staff of AVBS: Jonas Schneider, Julian Pudwell and Timo Kurzeia for their assistance during sampling campaigns.

