## Supplementary Material for "Treated wastewater irrigation promotes the spread of antibiotic resistance into subsoil pore-water"

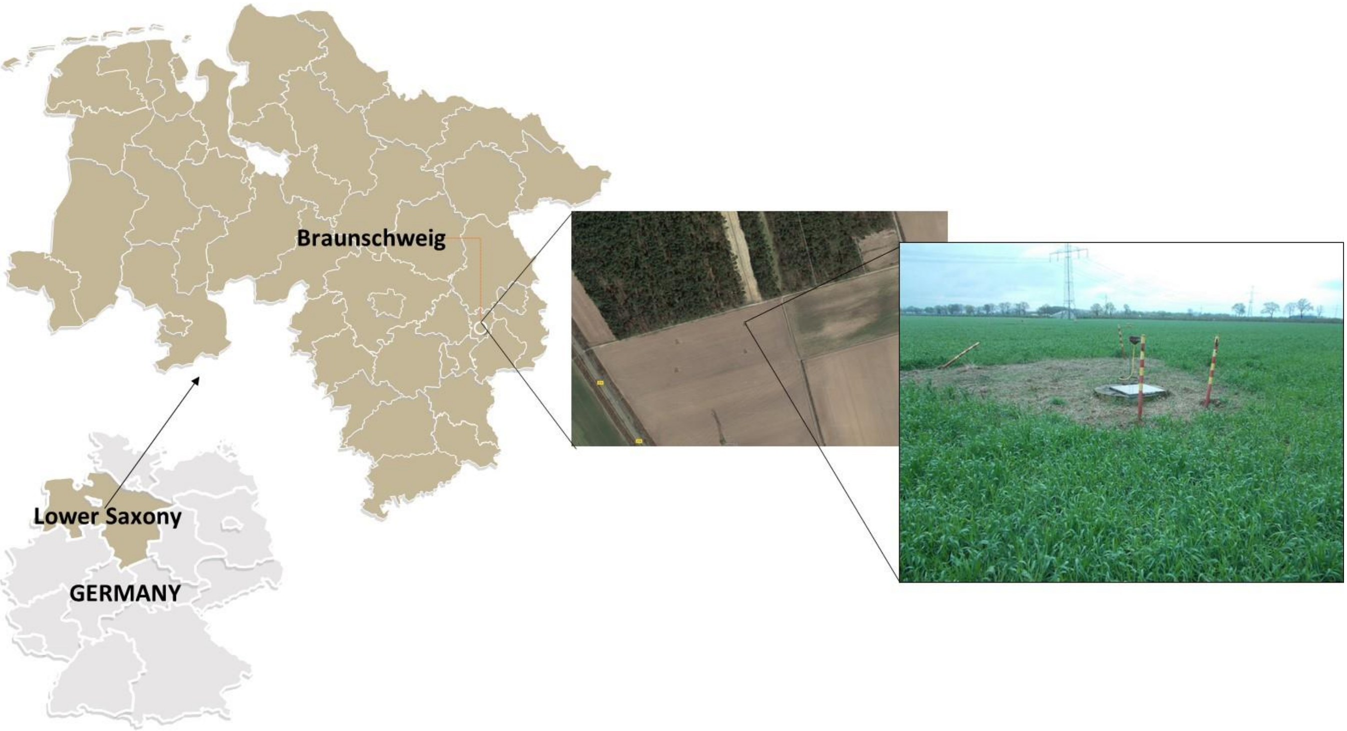


**Figure S1:** Field sampling location in the map. In the right picture, the entrance door to the Lysimeter installation.

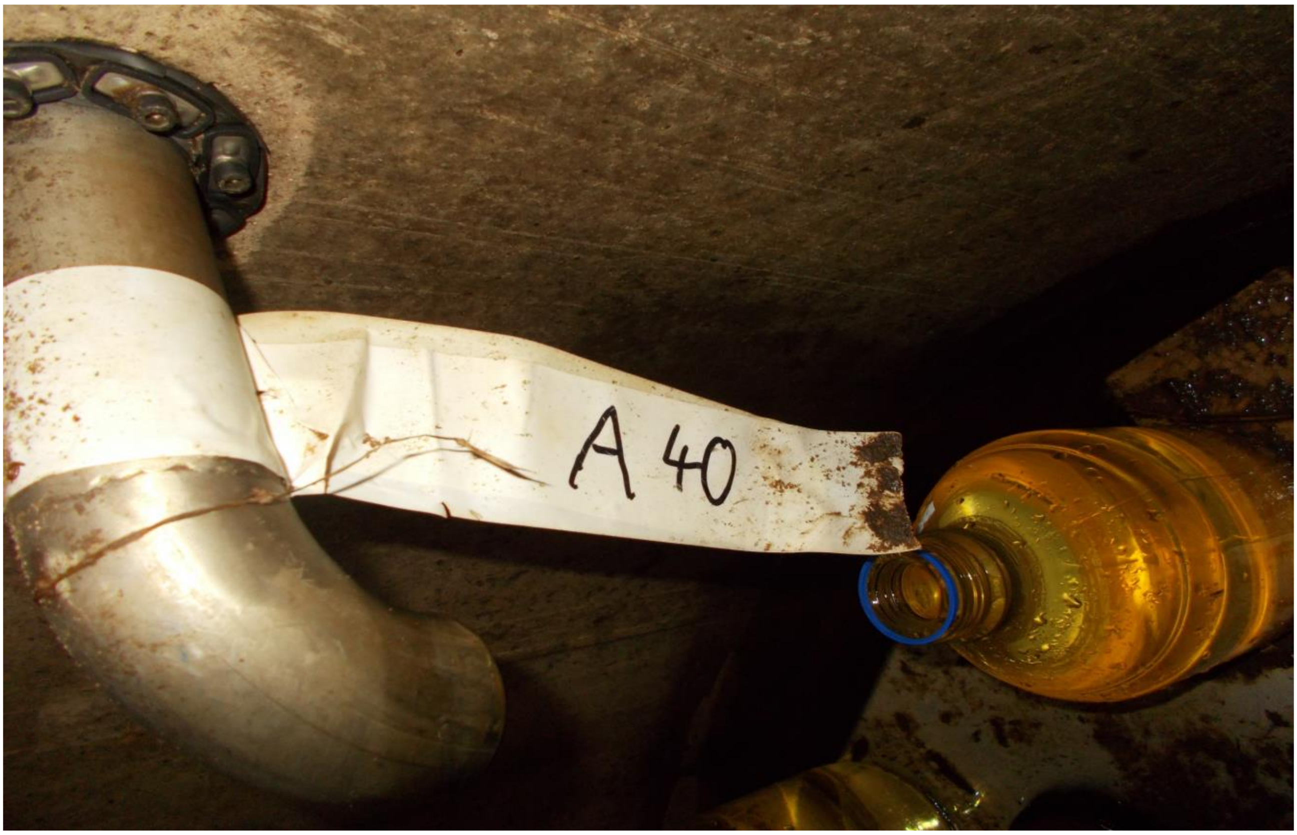

**Figure S2:** The interior bottom of the Lysimeter wells: a tap that allows collection of percolated water at a defined depth.

**Table S1:** Average annual chemical oxygen demand for the irrigation water provided by BWA (TWW & DS n= 36, TWW n= 60).

| | TWW | | TWW & DS | |
|---|---|---|---|---|
| | COD-homogenised (mg/L) | COD-filtrate (mg/L) | COD-homogenised (mg/L) | COD-filtrate (mg/L) |
| Maximum | 258 | 97 | 900 | 126 |
| Average | 62.8666667 | 44.5333333 | 468.25 | 96.5277778 |
| Minimum | 29 | 23 | 244 | 58 |

**Table S2:** Forest soil characteristics.

| **Characteristics** | **Forest Soil** |
|---|---|
| Clay (<2.0µm) % | 1.66 |
| Fine- Silt (2.0-6.3µm) % | 0.61 |
| Medium- Silt (6.3-20µm) % | 0.89 |
| Coarse-Silt (20-63µm) % | 1.69 |
| Fine-Sand (63-200µm) % | 20.2 |
| Medium-Sand (200-630µm) % | 61.5 |
| Coarse-Sand (630-2000µm) % | 11.3 |
| pH | 3.77 |
| Corg % | 4 |

**Table S3:** Tested genes, primers and protocols of qPCR assays.

| **Target gene** | | **Primers sequence** | **Amplicon size** | **Conditions** | **LOQ** | **Reference** |
|---|---|---|---|---|---|---|
| 16S rRNA (Indicator for microbial abundance) | Fw | TCCTACGGGAGGCAGCAGT | 195 bp | 95 °C - 10 min (1 cycle); 95 °C - 15 sec, 60 °C - 1 min (40 cycles) | 4000 copies per reaction | Cacace et al., 2019 |
| | Re | ATTACCGCGGCTGCTGG | | Other: 1 | | |

| | | | | | | |
|---|---|---|---|---|---|---|
| *bla*TEM (class A β-lactamase) | Fw | TTCCTG TTTTTG CTCACC CAG | 113 bp | 95 ºC - 10 min (1 cycle); 95 ºC - 15 sec, 60 ºC - 1 min (40 cycles) | 40-400 copies per reaction | Cacace et al., 2019 |
| | Re | CTCAAG GATCTT ACCGCT GTTG | | Other: 2 | | |
| *bla*CTX-M-32 (class A β- lactamase, cephalosporinase) | Fw | CGTCAC GCTGTT GTTAGG AA | 156 bp | 95 ºC - 10 min (1 cycle); 95 ºC - 15 sec, 58.5 ºC – 1 min (40 cycles) | 4-40 copies per reaction | Cacace et al., 2019 |
| | Re | CGCTCA TCAGCA CGATAA AG | | Other: 2 | | |
| *sul1* (sulphonamide resistant dihydropteroate synthase) | Fw | CGCACC GGAAAC ATCGCT GCAC | 162 bp | 95 ºC - 10 min (1 cycle); 95 ºC - 10 sec, 60 ºC -1 min (40 cycles) | 40-400 copies per reaction | Cacace et al., 2019 |
| | Re | TGAAGT TCCGCC GCAAGG CTCG | | Other: 2 | | |
| *qnrS* (protein family which protects DNA gyrase from the inhibition of quinolones) | Fw | GACGTG CTAACT TGCGTG | 118 bp | 95 ºC - 10 min (1 cycle); 95 ºC - 15 sec, 60 ºC - 1 min (40 cycles) | 4-40 copies per reaction | Cacace et al., 2019 |
| | Re | TGGCAT TGTTGG AAACTT | | Other: 2 | | |
| *intI1* (class I integrase, this gene is associated with horizontal gene transfer and environmental pollution) | Fw | GATCGG TCGAAT GCGTGT | 196 bp | 95 ºC - 10 min (1 cycle); 95 ºC - 15 sec, ºC - 1 min (40 cycles) | 40-400 copies per reaction | Cacace et al., 2019 |
| | Re | GCCTTG ATGTTA CCCGAG AG | | Other: 2 | | |
| *bla*OXA-58 (class D β-lactamase, carbapemenase) | Fw | CACTTA CAGGAA ACTTGG GGTCG | 79 bp | 95 ºC - 10 min (1 cycle); | 4-40 copies per reaction | Cacace et al., 2019 |

| | | | | | | |
|---|---|---|---|---|---|---|
| | | | | 95 ºC - 15 sec, 60 ºC - 1 min (40 cycles) | | |
| | Re | AGTGTGTTTAGAATGGTGATC | | Other: 2 | | |
| *tetM* (ribosomal protection protein that protects ribosome from the translation inhibition of tetracycline) | Fw | GCAATTCTACTGATTTCTGC | 186 bp | 95 ºC - 10 min (1 cycle); 95 ºC - 15 sec, 55 ºC - 1 min (40 cycles) | 40 copies per reaction | Cacace et al., 2019 |
| | Re | CTGTTTGATTACAATTTCCGC | | Other: 3 | | |

Other: 1* 0.5 µM primers, 2* 0.25 µM primers, 3* 0.2 µM primers and 0.1 mg/mL BSA.

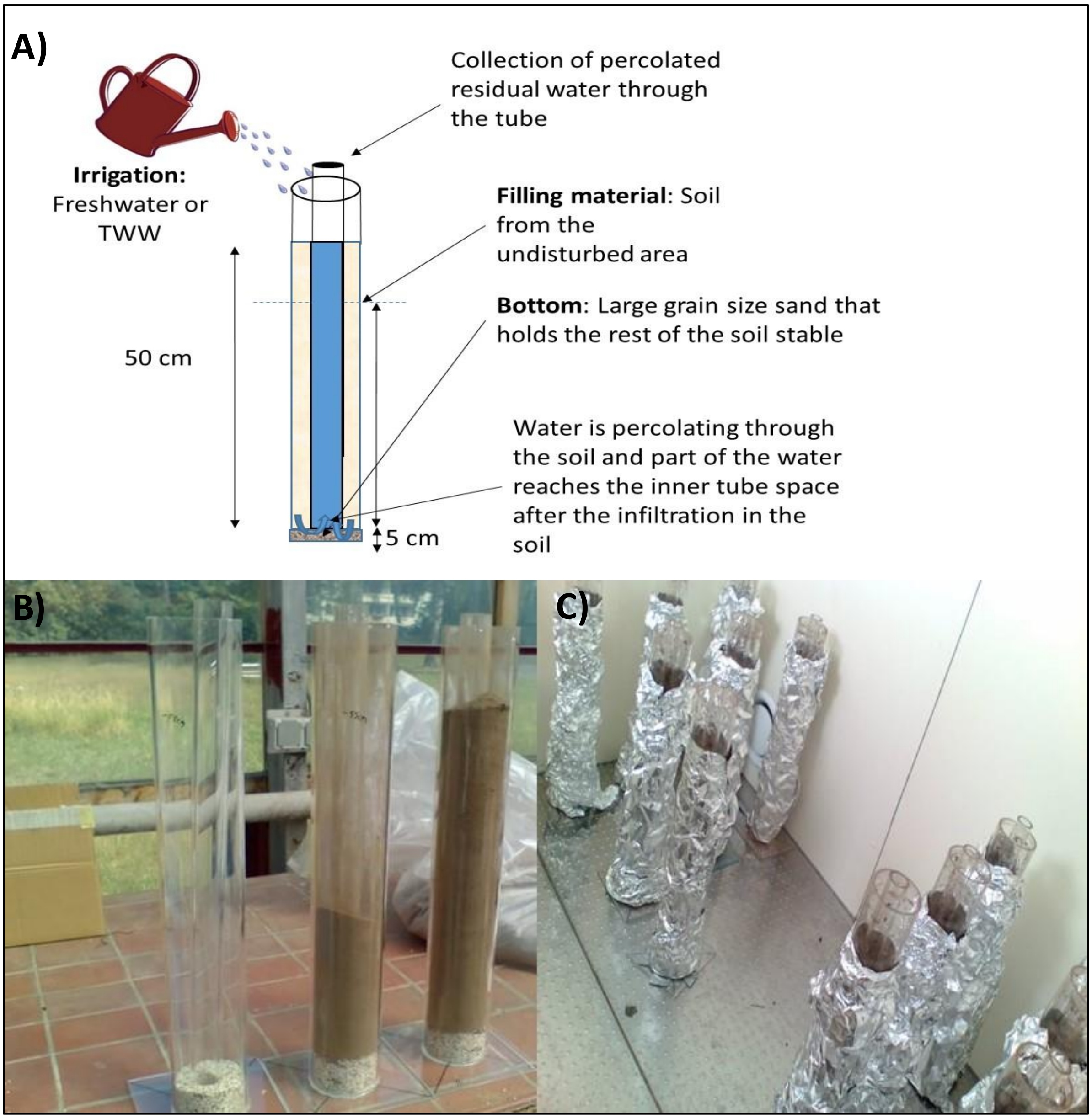


**Figure S3:** A) Schematic representation of the mesocosm design and structure, irrigation and percolated residual water sampling. B) The filling of the mesocosms with gravels in the bottom and sieved homogenized soil in the main tube. C) Aluminum foil wrapped mesocosms in the controlled-temperature room, with 20$^{o}$C stable temperature and 12 hours light per day.

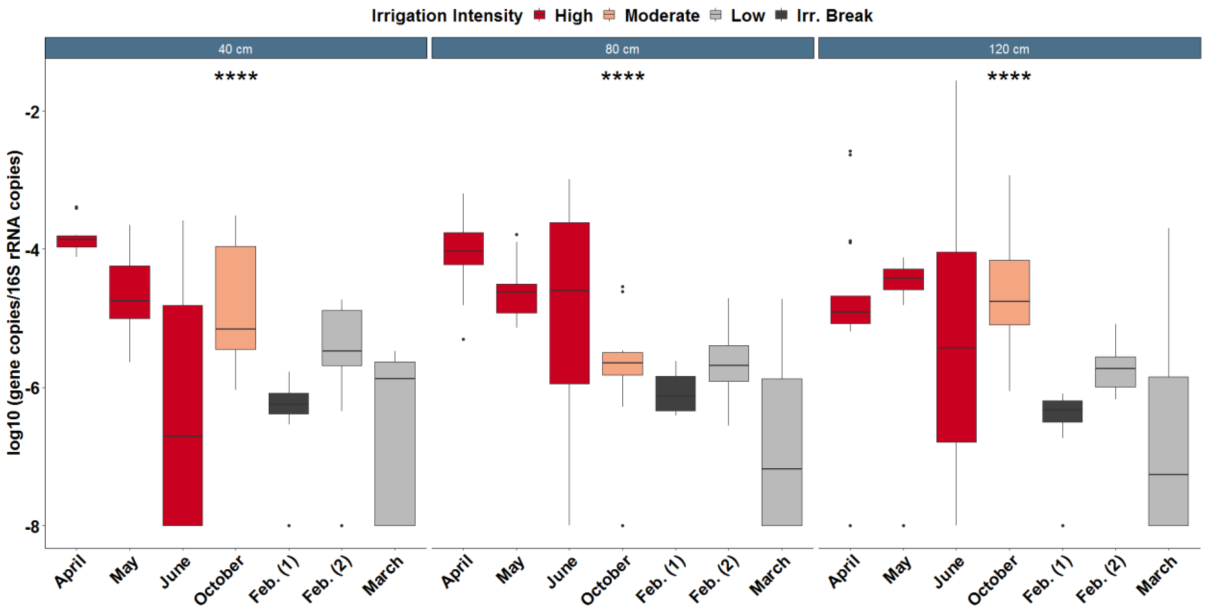


**Figure S4:** Relative abundance of *qnrS* during periods of high-, moderate-, low-irrigation intensity or irrigation break (Irr. Break). Kruskal-Wallis test: *p<0.05, **p<0.01, ***p< 0.001, ****p<0.0001, n=4.

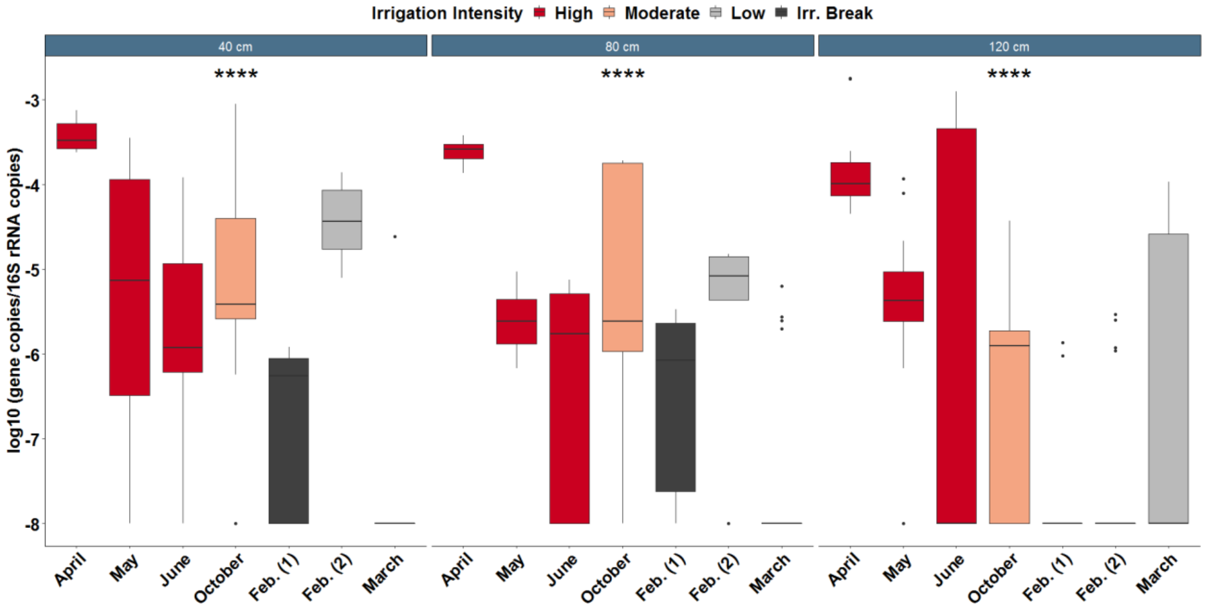


**Figure S5:** Relative abundance of *bla*$_{OXA-58}$ during periods of high-, moderate-, low-irrigation intensity or irrigation break (Irr. Break). Kruskal-Wallis test: *p<0.05, **p<0.01, ***p< 0.001, ****p<0.0001, n=4.

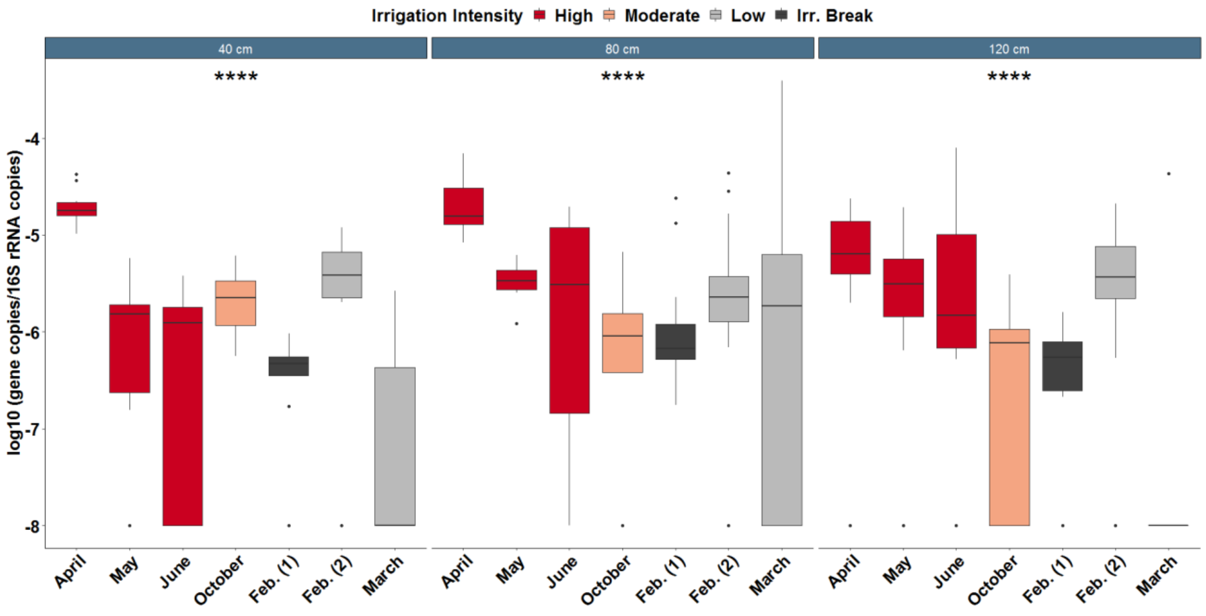


**Figure S6:** Relative abundance of *bla*CTX-M-32 during periods of high-, moderate-, low-irrigation intensity or irrigation break (Irr. Break). Kruskal-Wallis test: *p<0.05, **p<0.01, ***p< 0.001, ****p<0.0001, n=4.

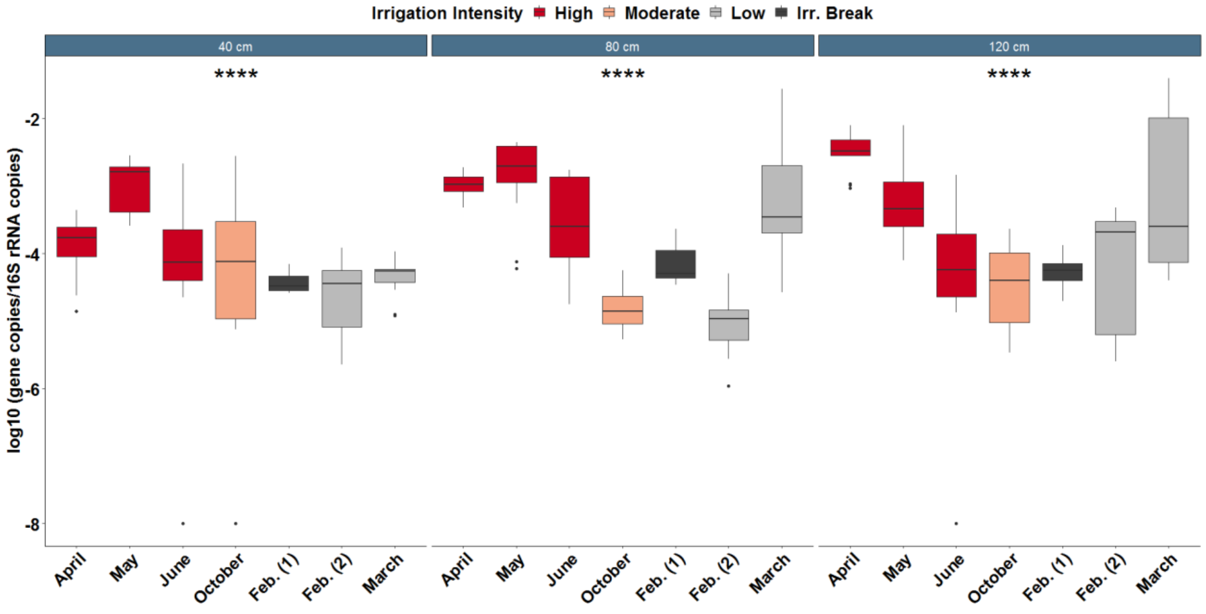


**Figure S7:** Relative abundance of *bla*TEM during periods of high-, moderate-, low-irrigation intensity or irrigation break (Irr. Break). Kruskal-Wallis test: *p<0.05, **p<0.01, ***p< 0.001, ****p<0.0001, n=4.

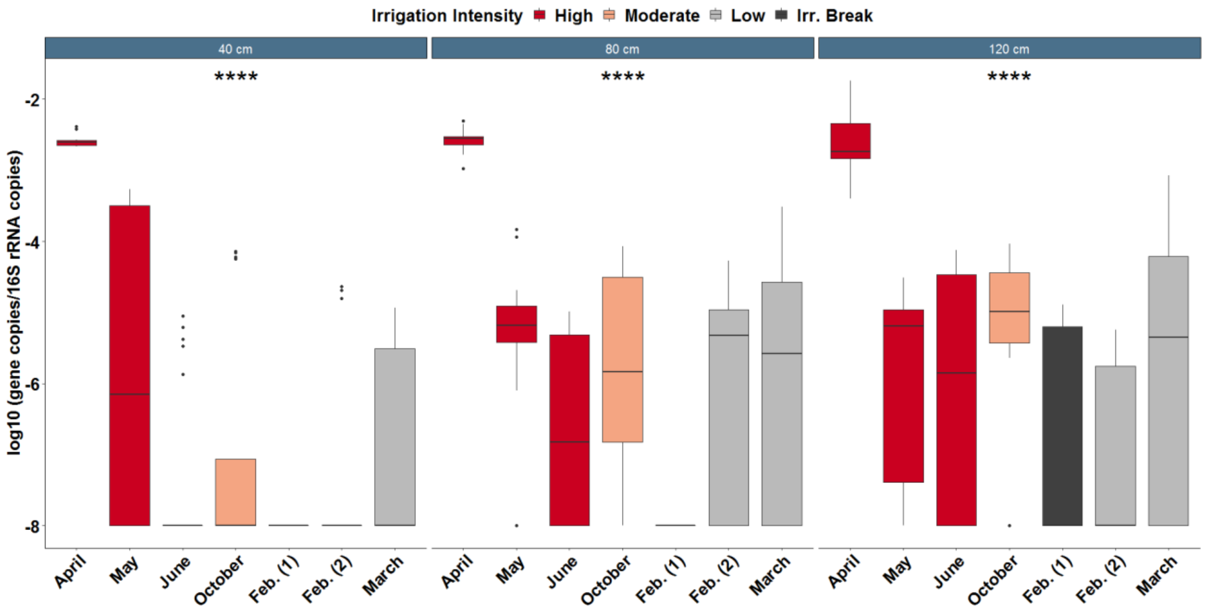


**Figure S8:** Relative abundance of *tetM* during periods of high-, moderate-, low-irrigation intensity or irrigation break (Irr. Break). Kruskal-Wallis test: *p<0.05, **p<0.01, ***p< 0.001, ****p<0.0001, n=4.

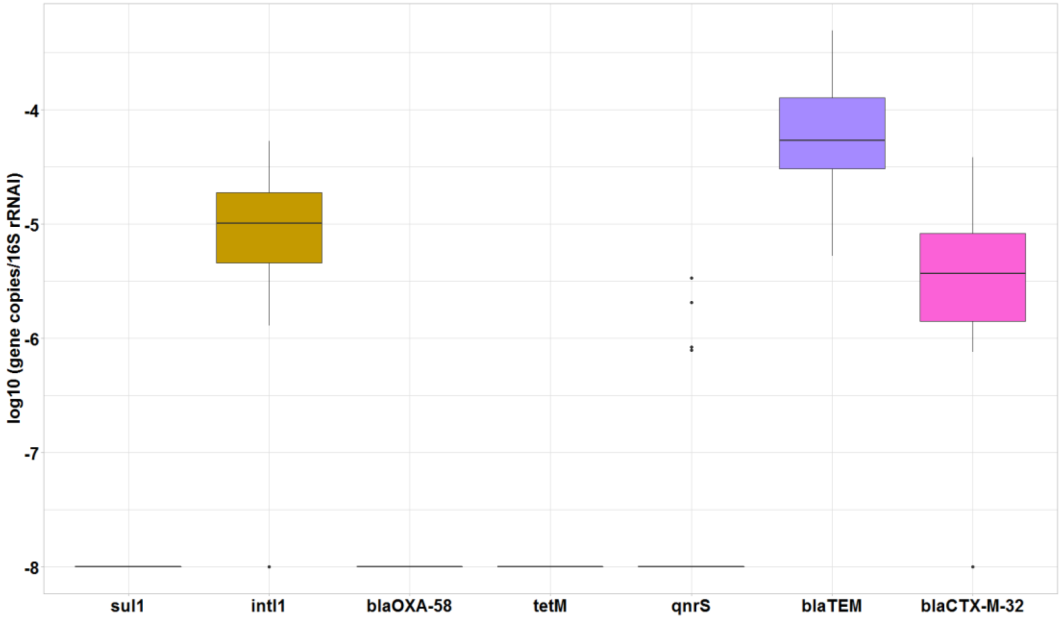


**Figure S9:** Relative abundance of ARGs and *intl1* in the soil of the forest, before it was used for mesocosms experiments.

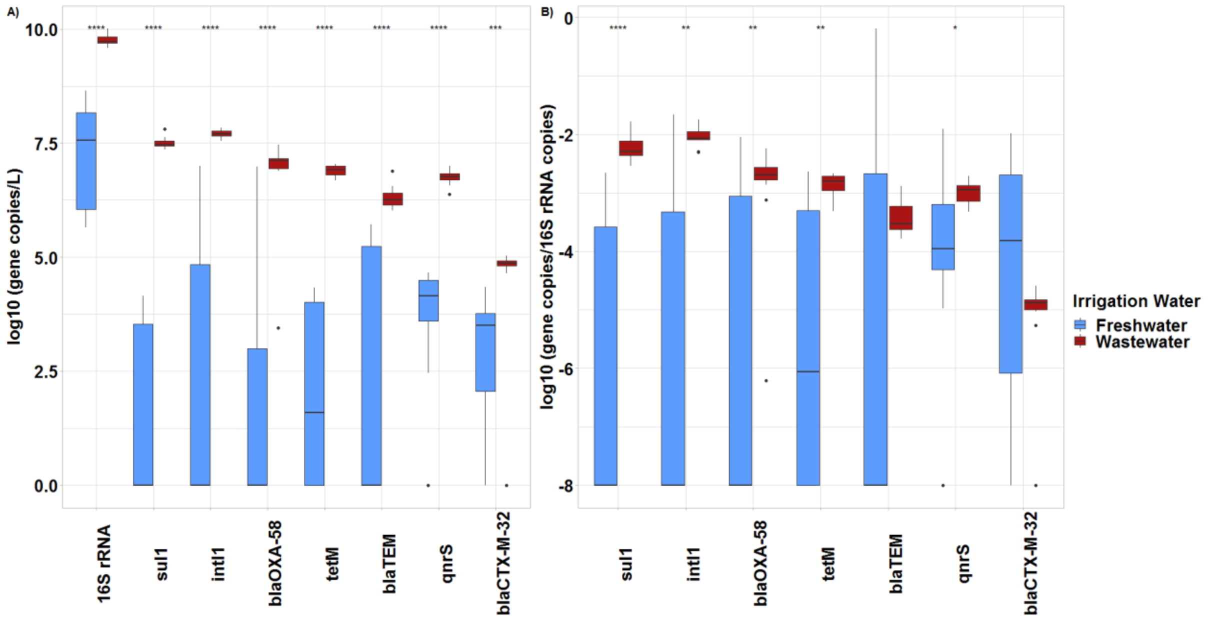


**Figure S10:** A) Absolute abundance of genes in copies/L and B) relative abundance of gene copies per 16S rRNA copy of the two types of irrigation water used for irrigation of the mesocosms (Wilcoxon rank sum test *p<0.05, **p<0.01, ***p< 0.001, ****p<0.0001, n=6).

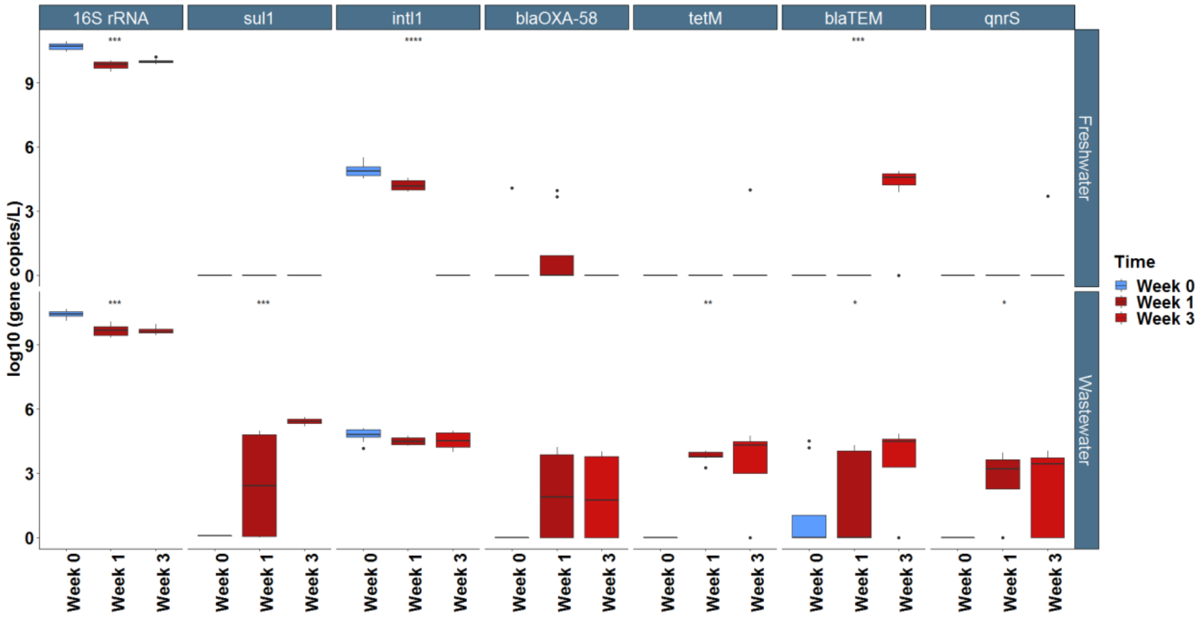


**Figure S11:** Absolute abundance of genes in the mesocosm percolated pore-water samples. (Kruskal-Wallis test, *p<0.05, **p<0.01, ***p< 0.001, ****p<0.0001, n=4).

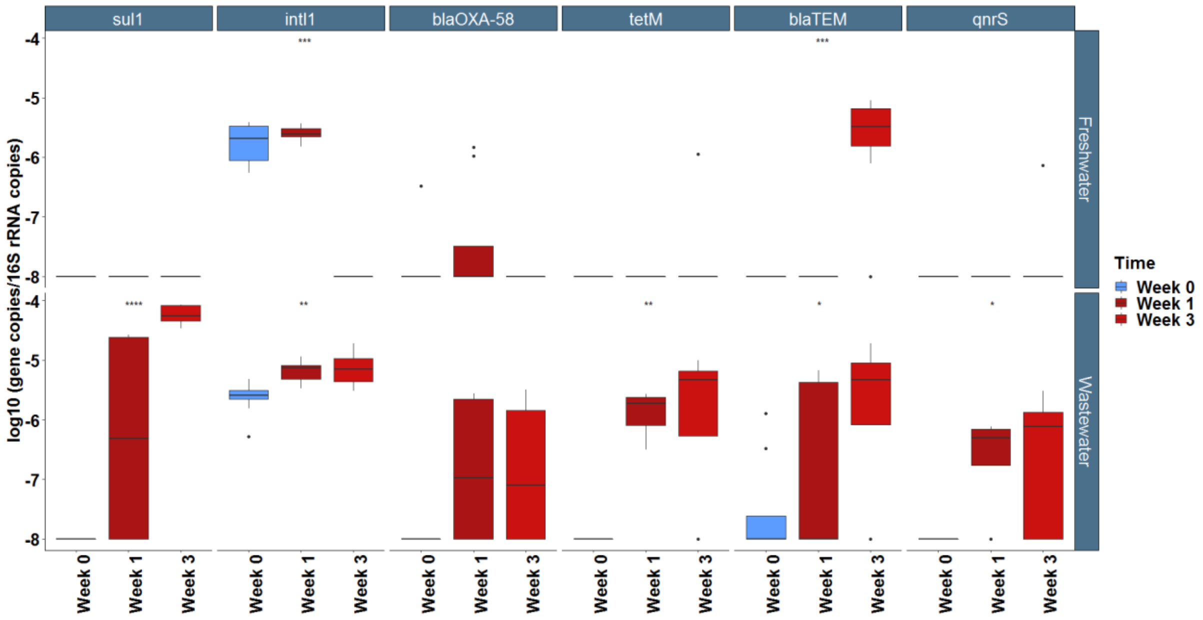


**Figure S12:** Relative abundance of genes in the mesocosm percolated pore-water samples. (Kruskal-Wallis test *p<0.05, **p<0.01, ***p< 0.001, ****p<0.0001, n=4).
